# Coherent theta oscillations in the cerebellum and supplementary motor area mediate visuomotor adaptation

**DOI:** 10.1101/2021.11.01.466768

**Authors:** Elinor Tzvi, Leila Gajiyeva, Laura Bindel, Gesa Hartwigsen, Joseph Classen

## Abstract

The cerebellum and its interaction with cortical areas play a key role in our ability to flexibly adapt a motor program in response to sensory input. Current knowledge about specific neural mechanisms underlying the process of visuomotor adaptation is however lacking. Using a novel placement of EEG electrodes to record electric activity from the cerebellum, we studied local cerebellar activity, as well as its coupling with neocortical activity to obtain direct neurophysiological markers of visuomotor adaptation in humans. We found increased theta (4-8Hz) power in “cerebellar” as well as cortical electrodes, when subjects first encountered a visual perturbation. Theta power decreased as subjects adapted to the perturbation, and rebounded when the perturbation was suddenly removed. This effect was observed in two distinct locations: a cerebellar cluster and a central cluster, which were localized in left cerebellar crus I (lCB) and right supplementary motor area (rSMA) using linear constrained minimum variance beamforming. Importantly, we found that better adaptation was associated with increased theta power in left cerebellar electrodes and a right sensorimotor cortex electrode. Finally, increased rSMA –> lCB connectivity was significantly decreased with adaptation. These results demonstrate that: (1) cerebellar theta power is markedly modulated over the course of visuomotor adaptation and (2) theta oscillations could serve as a key mechanism for communication within a cortico-cerebellar loop.

## Introduction

Adaptation to sudden environmental changes is at the root of basic human motor performance. For example, when walking on a moving train, our brain quickly produces an altered motor program using error-based learning mechanisms, that enables a steady and safe walking pattern. This process is referred to as sensorimotor adaptation. Previous behavioral studies in patients with cerebellar ataxia and cerebellar lesions due to stroke have demonstrated that the cerebellum plays a central role in this process (Criscimagna-Hemminger et al., 2010; Henriques et al., 2014; Schlerf et al., 2013; Tseng et al., 2007; Tzvi et al., 2021). Specifically, researchers hypothesized that sensory prediction errors, i.e. the difference between a predicted sensory outcome and the actual sensory feedback, is deficient in cerebellar patients, leading to worse performance in visuomotor adaptation tasks compared to healthy controls (Butcher et al., 2017; Henriques et al., 2014; Schlerf et al., 2013; Tseng et al., 2007; Wong et al., 2019). Others probed the role of the cerebellum in visuomotor adaptation using non-invasive electric stimulation, during or prior to task performance. Cerebellar direct current stimulation is thought to increase excitability in Purkinje cells, leading to inhibition of deep cerebellar nuclei and reduction of thalamic facilitation of cortical structures (Grimaldi et al., 2016). These studies showed that direct current stimulation of the cerebellar cortex led to enhanced visuomotor adaptation (Block and Celnik, 2013; Galea et al., 2011; Hardwick and Celnik, 2014; Weightman et al., 2020). Thus, enhanced performance in the visuomotor adaptation task might be driven by increased activity in the cerebellar cortex, leading to decreased connectivity with cortical areas. Note, however, that replication attempts of this effect were not successful (Jalali et al., 2017; Liew et al., 2018; Mamlins et al., 2019), probably due to strong inter-subject variability in task performance as well as in response to the stimulation. Imaging studies may overcome this limitation by investigating network interactions during performance in the visuomotor adaptation task. Indeed, these studies showed that the cerebellum and its interaction with cortical structures play an important role in visuomotor adaptation (Albert et al., 2009; Sami and Miall, 2013; Tzvi et al., 2020; Vahdat et al., 2011). However, functional MRI can only provide indirect information regarding neuronal mechanisms of sensorimotor adaptation.

The exact mechanism that allows flexible communication within a cortico-cerebellar network underlying visuomotor adaptation remains largely unknown. “Communication through coherence” has been previously suggested as a mechanism that allows interactions between neuronal populations through synchronous oscillations (Fries, 2005). Testing this hypothesis in humans would require recordings of neuronal oscillations from the cerebellum. However, although oscillations from the cerebellum can relatively easily be recorded in animals using local field potentials (LFP) (D’Angelo et al., 2001), it has been debated whether non-invasive recordings are possible in humans using electroencephalography (EEG) or magnetoencephalography (MEG) (Andersen et al., 2020). A recent modelling study of EEG/MEG signals, combined with high-resolution MRI of an ex-vivo human cerebellum, has nonetheless found that cerebellar signals could indeed be reliably captured from the surface of the head (Samuelsson et al., 2020). Furthermore, both evoked and oscillatory cerebellar activity were found using MEG, in response to omissions of tactile stimuli (Andersen and Dalal, 2021). Additionally, EEG studies found that high-frequency cerebellar oscillations (Todd et al., 2018a) as well as vestibular cerebellar evoked potentials (Todd et al., 2018b), could be reliably detected in electrodes placed over the posterior fossa in humans. Finally, Pan and colleagues (2020) demonstrated that pathological cerebellar oscillations related to essential tremor could be recorded non-invasively in human patients. Importantly, these oscillations were validated by congruent LFP signals recorded from cerebellar cortex in the animal model. These findings testify to the feasibility of recording cerebellar oscillations in humans using EEG. Therefore, we hypothesized that direct insight in the physiology of sensorimotor adaptation in humans could be obtained by cerebellar EEG.

In a recent fMRI study, we showed that cerebellar activity (along with other cortical structures), specifically increases as a response to a sudden exposure to a visuomotor rotation, decreases as visuomotor adaptation progresses, and increases again when the rotation is suddenly removed (Tzvi et al., 2020). In addition, directed connectivity analysis using Dynamic causal modelling revealed modulation of cerebellum to premotor cortex connectivity by adaptation. Moreover, modulation of this connection was associated with a faster return to the original routine. Building on these prior results, we set out to test whether sensorimotor adaptation exhibits: (1) a unique pattern of cortical and cerebellar oscillations and (2) coherent interactions between cerebellum and cortical structures. Our findings suggest that interactions within a cortico-cerebellar loop are mediated through coherent oscillations.

## 1. Materials and methods

### 1.1. Participants

26 healthy participants (mean age: 28 years, range 20-34; 7 males) took part in the experiment after giving informed consent. Participants were financially compensated for their participation. All participants were classified as right-handed by means of the Edinburgh Handedness Inventory (Oldfield, 1971) (score: 84 ± 15) and had normal or corrected to normal vision. Participants were non-smokers and did not suffer from any mental or neurologic disorder (by self-report). We excluded professional musicians and computer gamers. One subject was excluded due to an error in the data acquisition and another subject due to lack of adaptation (see section 2.1 below), resulting in a final sample of 24 subjects for the behavioral analysis. Two more subjects had to be excluded due to artifacts in the EEG signal and an error in electrode locations registration, resulting in a final cohort of 22 subjects for the EEG analyses. The study was approved by the Ethics Committee of the University of Leipzig (280/20-ek) and was performed in accordance with the Declaration of Helsinki.

### 1.2. Experimental paradigm and task design

During EEG recordings and task performance, subjects were seated comfortably in front of a 17” computer screen, about 0.75m away, on which visual stimuli were presented. The task was designed using Psychtoolbox-3 (Brainard, 1997) operating on MATLAB R2019b (Mathworks®). Prior and following task performance, we recorded EEG during resting-state (RS) for a duration of 200s each. Subjects performed a visuomotor adaptation task, previously described by (Galea et al., 2011), using a digital pen moved on a digital tablet (Wacom Intuos Pro L, Wacom, Kazo, Japan). The movements were visualized on the screen. A carton box covered the set-up to prevent visual feedback from the moving upper-limb. The position of the pen in 2D was sampled at 60Hz, which means that adjacent sampling points had 17ms between them. Subjects were instructed to perform straight, “shooting through” hand movements from a central starting point to a circular target in one of eight possible positions arrayed around a starting point at a distance of 70mm, equally distributed every 45° (Fig. 1A).

**Figure 1.**
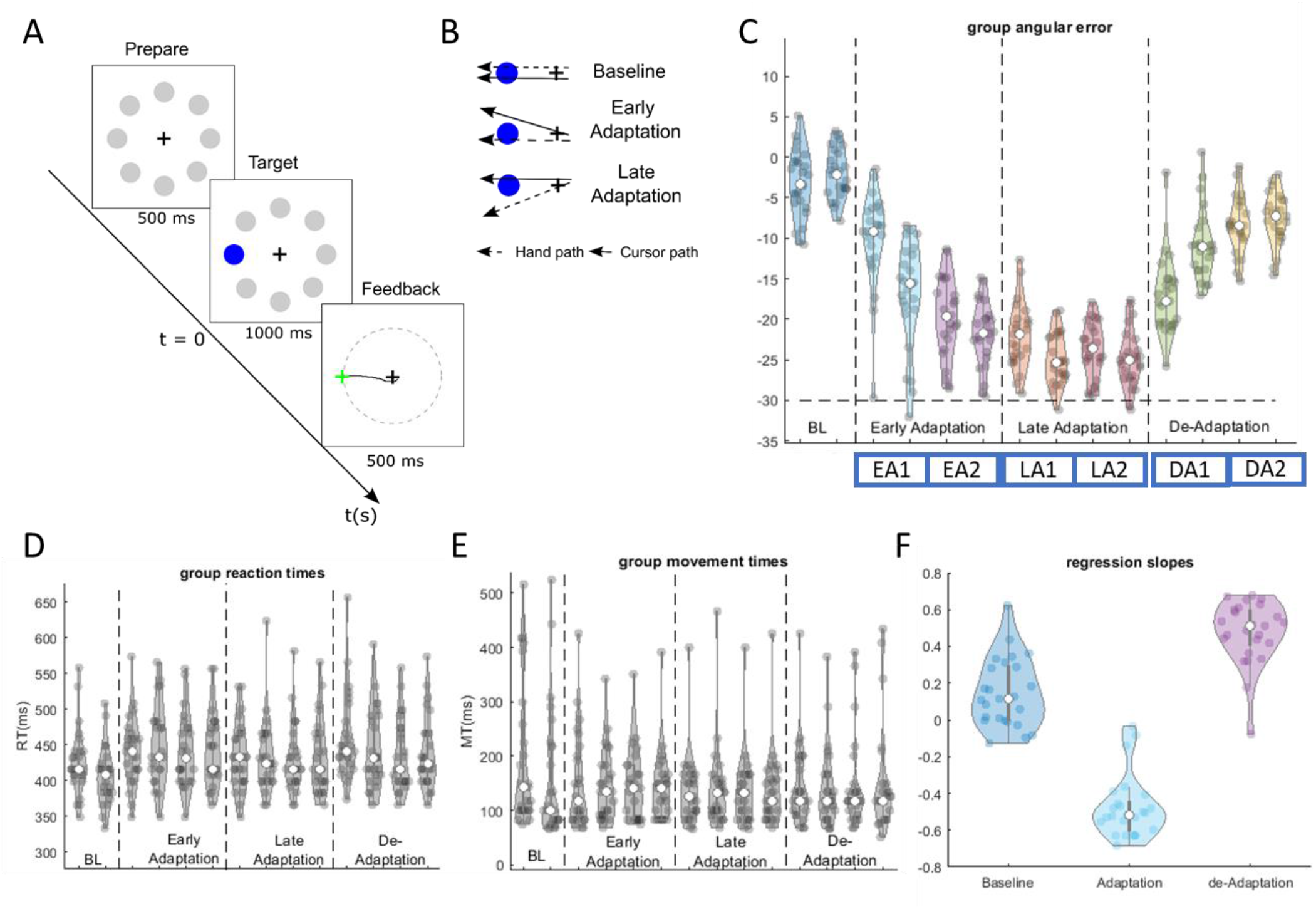
Experimental design and behavioral results. **A** Trial timeline. Each trial began with 500ms presentation of possible targets. Next, the current target was marked in blue, signaling the subjects to start the movement. This moment was defined as t = 0ms for the EEG analyses to follow. When subjects crossed the imaginary circle connecting the edges of all targets, all other stimuli disappeared, and feedback was given using a green cross at the crossing point. **B** Task conditions and the corresponding hand and cursor paths. **C** Violin plots for the angular errors across subjects. With time and across adaptation blocks (EA1, EA2, LA1, LA2), subjects reach the optimal value of −30°, representing their adaptation to a 30° visual shift in the cursor path. When the perturbation is removed (DA1, DA2), subject quickly re-adjust to the old routine. **D** Violin plots for reaction times across subjects, i.e., the time it took subjects to start the movement towards the target. **E** Violin plots for movement times across subjects, i.e., the interval between reaction time and the feedback. **E** Violinplots for linear regression slopes fitted for each subject.

The targets were presented pseudo-randomly such that every set of eight consecutive trials included one of each target positions. Trial description is presented in Fig. 1A. At the beginning of each trial, eight grey circles appeared around a center cross (i.e., the starting point) for 500ms. Next, the target was marked as a blue circle and cued the subjects to start the movement towards the target. Note that this point in time was defined as stimulus onset for the EEG analyses described below. Movement onset was defined to be the first time point in which more than 20-pixel deviance between adjacent sampling points (~17ms), in either direction, was detected. Once the pen crossed the invisible circle connecting the edges of all targets, all other stimuli disappeared, and a feedback was given as a green cross at movement end-point for 500ms (Fig. 1A). This time point was defined to be movement cessation. The cursor was visible at all times. To encourage fast performance, a bar at the bottom of the screen indicated the elapsed time.

Over the course of the experiment, a 30° clockwise visuomotor perturbation of the cursor movement on the screen, was introduced abruptly (Fig. 1B). Subjects were not aware of this manipulation prior to the experiment. Later in the experiment, this perturbation was removed, also abruptly. The experiment was therefore divided into three phases: baseline (BL) containing simple center-target movements without perturbation, adaption (ADP) containing a 30° clockwise perturbation of the cursor movement on the screen, and de-adaptation (DA) phase in which the perturbation was removed. The experiment consisted of a total of 14 task blocks, each with 24 trials. In between the blocks 10s breaks were introduced. Two blocks were included for the BL condition, the ADP and DA consisted of 8 and 4 blocks, respectively. The total task duration was ~20min, depending on individual performance.

Prior to the main experiment, subjects performed a short familiarization with the task which consisted eight trials without perturbation as well as eight trials with random visual perturbations between 30° counter clockwise and 30° clockwise. In contrast to the main experiment, subjects could see their upper extremities during the familiarization phase. Following the main experiment, we asked the subjects whether they noticed a visuomotor perturbation and to describe it. Their answers were rated according to the extent of explicit knowledge of the manipulation as well as the use of a cognitive strategy in the experiment (0 – no knowledge, 6 – full knowledge).

### 1.3. Behavioral analysis

Analysis of behavioral data was performed using custom-made scripts in MATLAB. We assessed learning in each trial using the angular error, defined as the angle between a line connecting the center cross and the target, and a line connecting the center cross and the position of the cursor at peak velocity. A negative angular error describes movements performed counter clockwise during ADP to counter-act the 30° clockwise visual perturbation and successfully hit the target. A positive angular error describes movements performed clockwise. We excluded trials that had an angular error > 60° in any direction. We then fitted a linear regression curve to the angular errors across all trials in conditions ADP and DA, in each subject. We used the regression slopes (Fig. 1F) to evaluate learning of the visuomotor perturbation as well as re-learning of the original mapping during DA. In addition, we calculated the median angular error in each block (across 24 trials).

Reaction times (RT) and movement times (MT) were also analyzed (Fig. 1D-E). RT was defined as the time difference between the appearance of a target circle (circle color changes from grey to blue, Fig. 1A) and movement onset (defined above). MT was defined as the duration between movement onset and movement cessation. Outliers were excluded as described above. We calculated the median RT and MT in each block for each subject.

### 1.4. MRI recordings

MRI images were recorded using a 3T Siemens Skyra or Siemens Prisma head-scanner with 32-channel head coil at the Max Planck Institute for Human Cognitive and Brain Sciences in Leipzig, Germany. A high resolution T1-weighted gradient-echo structural image was acquired using the following parameters: image matrix: 240 x 256, 176 sagittal slices of 1mm thickness, TR = 2300ms, TE = 2.98ms.

### 1.5. EEG recordings

EEG was recorded using Ag/AgCl electrodes embedded in a custom 64-channel cap and connected to an eego™ amplifier (ANT Neuro b.v., Hengelo, the Netherlands) with a sampling-rate of 512Hz and 24bit resolution. A low-pass filter was applied at 0.26*sampling-rate (~133Hz). Electrodes were placed according to an extension of the 10-20 system at the following locations: Fp1, Fpz, Fp2, AFz (ground), F7, F5, F3, F1, Fz, F2, F4, F6, F8, FC5, FC3, FC1, FCz, FC2, FC4, FC6, T7, C5, C3, C1, Cz, C2, C4, C6, T8, TP7, CP5, CP3, CP1, CPz (online reference), CP2, CP4, CP6, TP8, P7, P5, P3, P1, Pz, P2, P4, P6, P8, PO7, PO5, PO3, POz, PO4, PO6, PO8, O1, Oz, O2, Iz. Six additional locations were recorded: CB11, CB1z, CB12, CB21, CB2z, CB22, located 5% inferior to Iz, and 10% laterally distributed over both left and right cerebellar hemispheres (yellow electrodes in Fig. 2A). Electrode impedances were kept below 5kΩ.

**Figure 2.**
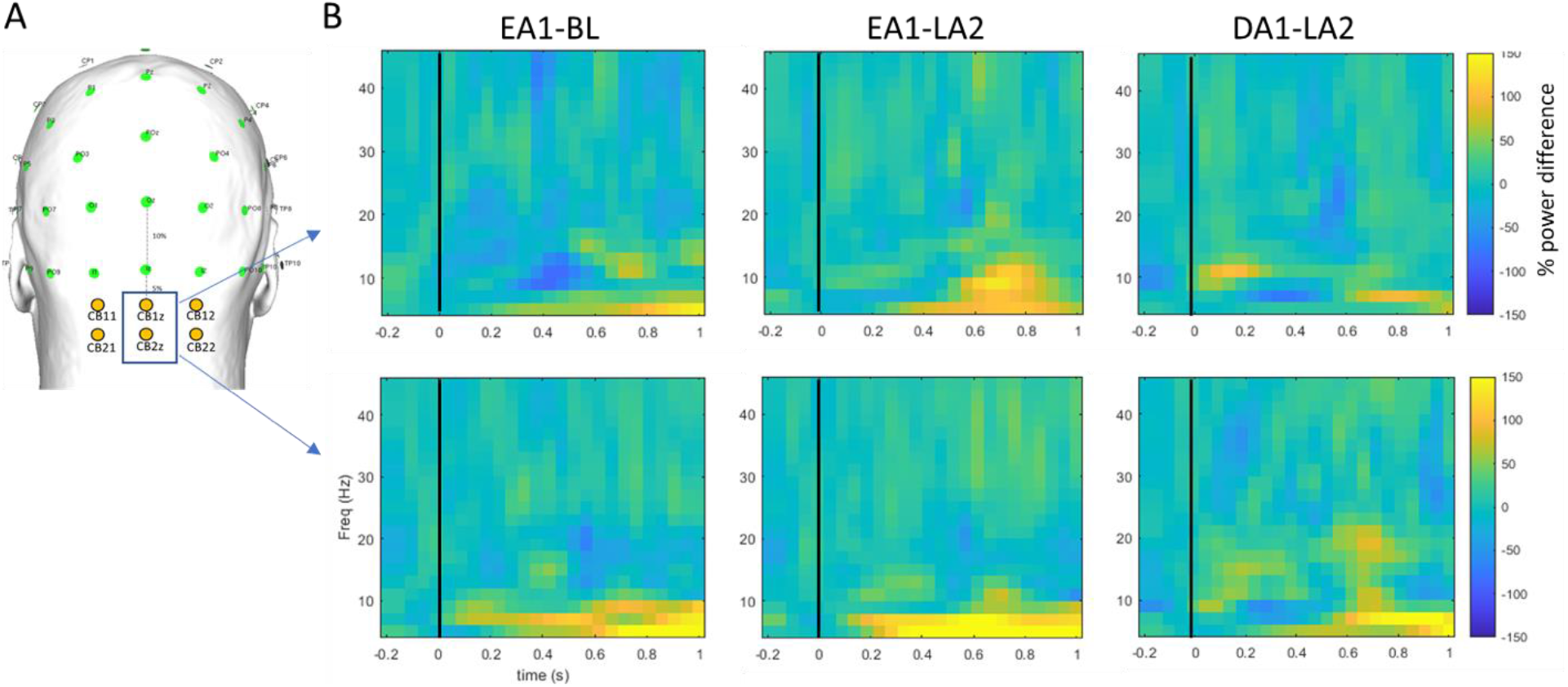
**A** Locations of cerebellar electrodes (in yellow) on the surface of the head. **B** Time-frequency representations for locations CB1z and CB2z for three condition differences: early adaptation (EA1)-baseline (BL), early adaptation (EA1)-late adaptation (LA2), and de-adaptation (DA1)-late adaptation (LA2).

### 1.6. EEG pre-processing

Pre-processing and all subsequent analyses were performed using in-house Matlab scripts, EEGLAB toolbox (Delorme and Makeig, 2004) and Fieldtrip toolbox (Oostenveld et al., 2010). We first applied a high-order finite-impulse response band-pass filter (F_*cutoff*_ = 1-49Hz) to remove slow drifts and power line noise. Signals were then re-referenced offline to the average of the signal from left and right mastoids and the signal from electrode CPz was re-calculated. Then, signals were segmented into 3s epochs (For task-based signals: −1s to 2s around stimulus onset). An independent component analysis (ICA) procedure was then applied to the signals to identify components related to eye blinks and horizontal eye movements based on their topography and signal shape (Makeig et al., 2004). In most subjects we removed 2-4 components. Additional artifacts were removed using a simple threshold (−70µV, +70µV) on the filtered data.

### 1.7. Spectral power and statistical analysis

Next, we computed the power spectrum of the EEG signals using the Morlet wavelet. Signals were filtered to obtain oscillatory power at 1-49Hz using wavelets of seven cycle length. Frequency resolution was set to 2Hz and time resolution to 50ms. Task-based signals were then averaged across a 0–500ms time window (0ms being stimulus onset), which mostly accounts for motor preparatory signals (“PREP”), and across a 500–1000ms time window, which account for both the movement and the feedback (“MOV”). Note that since the movement itself was very short, it was not possible to disentangle these processes. Resting-state (RS) signals were averaged across the entire time window. Next and in order to limit the number of comparisons, we averaged the signals across the different frequency bands of interest: theta (4-8 Hz), alpha (9-13Hz), beta (14-30Hz), gamma (31-49Hz). No baseline correction was applied. To enhance signal-to-noise ratio in task-based signals, we concatenated two blocks of 24 trials each for the EEG analysis resulting in one block for BL, four blocks for ADP: two early adaptation blocks (EA1, EA2) and two late adaptation blocks (LA1, LA2) as well as two blocks for DA (DA1, DA2).

Statistical tests of task-based signals were performed using ‘ft_statfun_depsamplesFunivariate’ (for rmANOVA) across the four ADP blocks (EA1, EA2, LA1, LA2). Statistical tests of resting-state signals were performed using ‘ft_statfun_depsamplesT’ to examine changes in pre-task (“PRE”) vs. post-task (“POST”) resting-state EEG. Both analyses employed Monte Carlo permutation testing with 1000 randomizations. Clusters were specified using the Fieldtrip function ‘ft_prepare_neighbours’ as channel neighbors with a distance smaller than 40mm. On average, for each channel, 4.1 neighboring channels were specified. Cerebellar channels were defined as a sperate cluster without any neighbors.

Spectral power changes were assessed in the different time windows (PREP: 0 - 500 ms, MOV: 500 - 1000 ms), in the different frequency bands (theta, alpha, beta, gamma). We report the cluster statistics as the sum of t-values for each electrode in the cluster, and the corrected p-value for the entire cluster. Significant clusters were defined based on a p-level of 0.05. Post-hoc Wilcoxon signed-rank tests were performed across all electrodes in the identified cluster of the main analysis. P-values were corrected using the false discovery rate (FDR).

### 1.8. Source analysis

For source reconstruction, we used the linear constrained minimum variance (LCMV) beamforming approach. As a first step we created a head model, specifying the individual head geometry based on T1 images of each subject (see specifications above), as well as tissue conduction properties, and applied the boundary element method (BEM) as a forward model. To this end, we segmented the T1 image into three tissue types: brain, skull and scalp and estimated for each tissue type a boundary triangle mesh (brain: 3000 points, skull: 2000 points and scalp: 1000 points). We then aligned the individual electrode positions, recorded using Localite EEG PinPoint (Localite GmbH, Germany), to the head model. A source model, specifying the locations of sources within the head model was then constructed as a discrete grid based on the head model geometry. For each grid point, a lead field matrix was calculated, which was subsequently used to calculate the inverse spatial filter using LCMV. Following the pre-processing steps described above, signals were re-referenced to a common-average reference and band-passed filtered (FIR, 4th order) to extract theta (4-8Hz) oscillations in a time window of 500–1000ms (stimulus onset: 0ms), based on the findings from the electrode-space analysis (see section 2.2. below). Source orientation was optimized by using the orientation of maximum signal power. To allow group level comparisons, individual source signals were then normalized using an MNI template. Statistical analyses of the source data were performed using cluster-level Monte Carlo permutation testing. Significant clusters were defined based on a p-level of 0.05.

### 1.9. Connectivity analysis

To study causal interactions between regions, we computed the phase-slope index (PSI, Nolte et al., 2008):

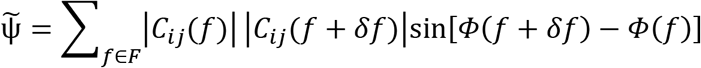

Where, 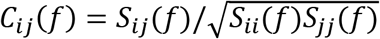 is the complex coherency, S is the cross-spectral matrix, *δf* is the frequency resolution, *ϕ* is the phase spectrum and F is the set of frequencies over which the slope is summed. 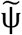 is then normalized by an estimate of the standard deviation: 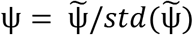.

The PSI is a highly robust measure of connectivity between EEG signals which serves to estimate the direction of information flux. Thus, a positive PSI indicates that the signal from the first region is leading the signal from the second region and vice versa. To this end, we first computed the signals from the two identified ROIs in source space showing significant power changes with adaptation, namely: left cerebellar crus I (lCB) ROI and right supplementary motor area (rSMA) ROI (see section 2.2 below). To this end, we used the peak activity and individual spatial filters found in the source localization analysis described above. Then, spectral representations of the reconstructed signals were obtained using the fast Fourier transform (FFT) and multi-tapers (N = 5). Next, we evaluated PSI using the Fieldtrip function ‘ft_connectivity_psi’ in each condition and block. Statistical analysis was performed using Wilcoxon signed-rank tests, comparing connectivity across the different conditions.

## 2. Results

### 2.1. Behavioral analysis

One subject was excluded from the behavioral analyses due to extreme small slopes in both adaptation (ADP) and de-adaptation (DA) blocks compared to the group’s median, as well as extreme slow reaction-times compared to the group’s median. Individual boxplots for the angular error in each block are presented in supp. Fig. 1. Three subjects were fully aware of the visuomotor perturbation. As this explicit recognition could have developed during the task, these subjects were not excluded from further analyses.

Differences between regressions slopes in BL and ADP as well as between ADP and DA and between BL and DA were highly significant (p < e-12, p < e-15 and p < e-6, resp.). Violin plots for the group angular error in each block are presented in Fig. 1C. A linear regression curve was fitted for each subject for baseline (BL), ADP and DA. The negative slope of the regression in ADP indicates adaptation to the visuomotor perturbation. The positive slope of the regression in DA indicates successful de-adaptation or wash-out of the visuomotor perturbation, and return to the original visuomotor routine.

Reaction times analysis showed that subjects were slower in initiating movements to the visual target when the perturbation was introduced in the first block of ADP (compared to last block of BL, Z = 3.7, p < 0.001, Fig. 1D) as well as when the perturbation was removed in the first block of DA (compared to last block of ADP, Z = 3.1, p = 0.002, Fig. 1D). There were no differences between the first and the last block of ADP (p = 0.4), suggesting that subjects did not improve on movement initiation during ADP. Movement times (Fig. 1E) did not differ between conditions and blocks (p > 0.1).

### 2.2. Cerebellar oscillations underlie visuomotor adaptation dynamics

Next, we examined changes in cerebellar oscillatory power associated with visuomotor adaptation. To this end, we submitted power values in the different frequency bands (theta - θ, alpha - α, beta - β, gamma - γ), the two time-windows (PREP: 0 - 500ms, MOV: 500 - 1000ms) and across ADP blocks (EA1, EA2, LA1, LA2) to a rmANOVA and cluster-based Monte-carlo permutation analysis. We found a significant cluster (clusterstat = 96.1, p < 0.001) showing effects in both time-windows and across all frequency bands (see supp. Table 1 for an overview on the specific cerebellar locations and frequency bands showing significant effects). While these effects are meaningful for visuomotor adaptation, they might be driven by non-specific changes in the oscillatory signal with time. To gain further insight in processes that are associated with visuomotor adaptation, we additionally investigated comparisons of the different adaptation phases to baseline (BL) and de-adaptation (DA).

In Fig. 2B, we present exemplary time-frequency representations in electrodes CB1z and CB2z, averaged across the subjects at: (1) early adaptation compared to baseline (EA1-BL), (2) early adaptation compared to late adaptation (EA1-LA2) and (3) de-adaptation compared to the late adaptation (DA1-LA2). Increased θ power was evident across all the different comparisons, around 500–1000 ms following stimulus onset (appearance of the blue target circle), predominantly in electrode CB2z. Note that group reaction times were around 400-450 ms, suggesting that these effects on θ power occur during movement to the target.

Based on these time-frequency representations of changes across adaptation phases, we followed the statistical analysis above with post-hoc tests in the different frequency bands and the different time windows separately. Specifically, we compared EA1 to BL to tap into oscillatory activity associated with the first exposure to the visual perturbation, as well as compared DA1 to LA2, i.e., when the perturbation was suddenly removed.

Indeed, we found a significant cerebellar cluster (electrodes: CB11, CB21, CB2z, p = 0.02, clusterstat = 6.8) in MOV time-window, showing increased θ power comparing EA1 to BL. There were no significant clusters for PREP. In source space (data not shown), we identified a peak activation in left cerebellum crus I (peak voxel: t_21_ = 3.1, p = 0.006), left inferior occipital lobe (peak voxel: t_21_ = 3.3, p = 0.002) as well as right cerebellar lobule VI (peak voxel: t_21_ = 3.1, p = 0.002) for the comparison between EA1 and BL. θ power then decreased with adaptation in all cerebellar electrodes as revealed by post-hoc Wilcoxon signed-rank tests showing significant effects comparing EA1 to LA2 (all Z > 2.9, p < 0.004, FDR corrected, Fig. 3B).

**Figure 3.**
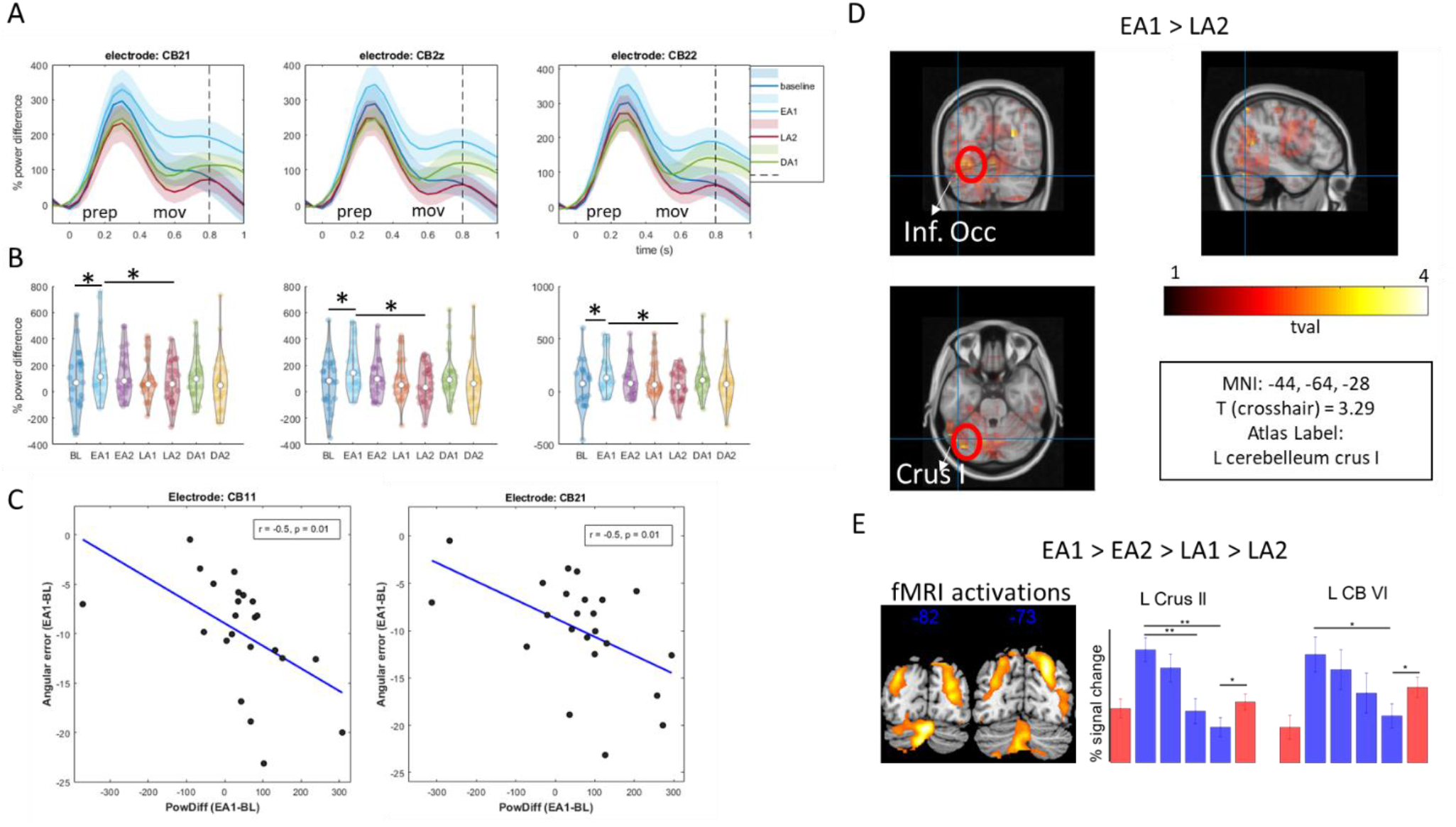
**A** Theta oscillations in cerebellar electrodes across the different conditions. Specific for the MOV time window, theta power (% difference compared to pre-stimulus baseline) increased at early adaptation (EA1) compared to baseline (BL), attenuated as adaptation progressed, and rebounded at de-adaptation (DA1). Shaded areas are standard deviations of the mean across subjects. Dashed lines indicate the time point for which power values were extracted for B. **B** Violinplots for t = 0.8s, at the peak of the MOV time window. Significant differences between conditions are marked with an asterisk for illustration. **C** Negative correlations between angular error and theta power difference in EA1-BL suggest that subjects that could better adapt early on, also exhibit larger theta power difference. **D** Source reconstruction for EA1-BL effects. **E** Activation patterns from previous fMRI study (Tzvi et al, Neuroimage 2020), showing differences from EA1 to LA2 and the corresponding pattern extracted from left cerebellar locations.

Reconstruction of this θ power changes in source space revealed peak activity in left cerebellar crus I (peak voxel: t_21_ = 3.3, p = 0.002, Fig. 3D) as well as left inferior occipital lobe (peak voxel: t_21_ = 3.7, p = 0.002, Fig. 3D). There were no significant activation clusters in right cerebellum or right occipital cortex. Comparing DA1 to LA2, we found a trend for θ power increase in cerebellar electrode CB22 (Z = 2.1, p = 0.03, uncorrected). θ power was then significantly decreased from DA1 to DA2 in electrode CB22 (Z = 2.8, p = 0.005, FDR corrected). Here, source reconstruction revealed a small activation cluster in right cerebellum crus II (peak voxel: t_21_ = 4.1, p = 0.002, data not shown).

In addition, we found significant α power increase in MOV (electrodes: CB21, CB22, p = 0.01, clusterstat = 4.7, data not shown) when comparing DA1 to LA2. α power did not change from EA1 to LA2 or between EA1 and BL (all p > 0.05). Analyses of PREP time-window and of β and γ power yielded non-significant effects using post-hoc tests (p > 0.05). We therefore focused the next analyses on θ power and MOV time-window.

Next, we extracted EA1-BL θ power differences in MOV and electrodes CB11, CB21 and CB2z (showing significant EA1-BL power difference) and examined their association with differences in angular errors in EA1-BL. Here we focused on a specific time point in which maximal θ power was evident (t = 800ms following stimulus onset, see Fig. 3B). We found a significant negative correlation in left cerebellar electrodes CB11 and CB21 (r = −0.52, - 0.51 resp., p = 0.01, FDR corrected, Fig. 3C). This suggests that the observed θ power increase in EA1 was associated with stronger adaptation to the visuomotor rotation early-on. There were no correlations between power differences in other conditions and angular errors.

### 2.3. Centro-parietal theta oscillations underlie visuomotor adaptation dynamics

We next explored the effect of visuomotor adaptation on oscillations in cortical locations by performing the exact same analysis as above (rmANOVA across both time windows, frequency bands, and ADP blocks) in all other electrodes. We found three significant clusters: a centro-parietal θ cluster, a frontal α cluster and a wide-spread β cluster, showing effects in both time-windows (see supp. Table 1 for an overview on the specific cortical locations). However, to limit the number of statistical comparisons and specifically associate the effects found in the cerebellar electrodes above, we focused the following analyses on θ and the MOV time window.

First, we identified a central cluster (p = 0.01, clusterstat = 34.9, Fig. 4A) showing decreased θ power with adaptation. Post-hoc Wilcoxon signed-rank tests showed that θ power was significantly larger comparing EA1 to LA2 (all Z > 3.0, all p < 0.003, FDR corrected). Source reconstruction of this effect revealed a peak activity in right supplementary motor area (rSMA, peak voxel: t_21_ = 3.5, p = 0.002, Fig. 4C). θ power significantly increased from LA2 to DA1 in electrodes CP1, CP2, and CPz (all Z > 2.4, p < 0.02, FDR corrected). This θ power difference was associated with a source in right SMA (peak voxel: t_21_ = 3.3, p = 0.002).

**Figure 4.**
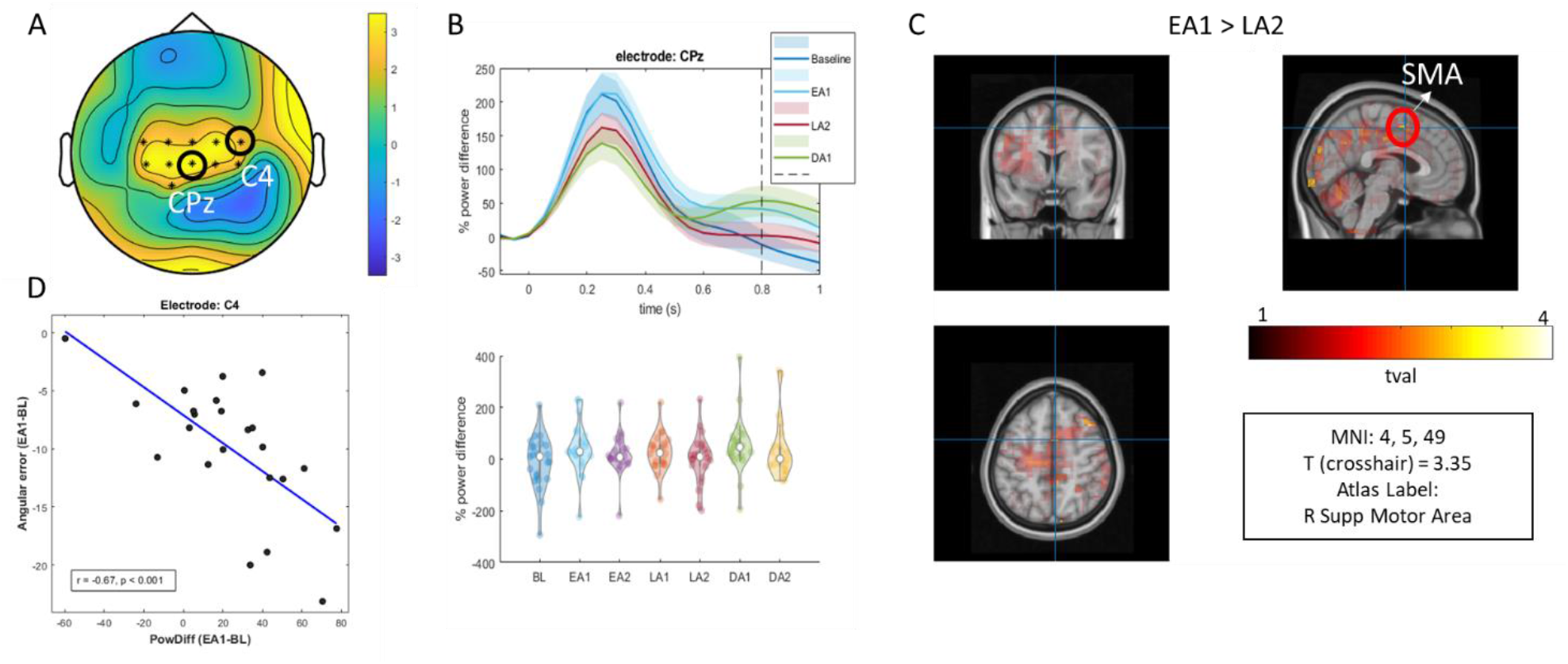
**A** Topographic plot showing a cluster (locations marked with an asterisk) of significant theta power changes across adaptation. **B** Theta oscillations in electrode CPz. **C** Source reconstruction of EA1-LA2 theta power. **D** Negative correlation between theta power and angular errors in EA1-BL suggest that subjects that could better adapt early on, also exhibit larger theta power difference in C4.

We next examined an association between EA1-BL θ power differences and EA1-BL differences in angular error, focusing on t = 800 ms following stimulus onset, similar to the analysis in cerebellar electrodes above. In electrode C4, we found a significant negative correlation (r = −0.67, p < 0.001, FDR corrected, Fig. 4D), suggesting that similar to the left cerebellar effects above, θ power increase in EA1 was associated with stronger adaptation to the visuomotor rotation early-on. There were no correlations between power differences in other conditions and behavior.

### 2.4. Modulation of resting-state oscillations by visuomotor adaptation

Next, we also analyzed possible changes in resting-state (RS) cerebellar oscillatory power following visuomotor adaptation, by examining the difference between pre-task RS (PRE) and post-task RS (POST) across the different frequency bands (θ, α, β, γ). We found a significant cluster (clusterstat = 13.19, p = 0.019) specific for θ, in all cerebellar electrodes except CB12, showing increased θ power from PRE to POST (Fig. 5A). Across cortical locations using the same analysis in all other electrodes, we found a significant cluster (clusterstat = 318, p < 0.001) showing increased θ but also α and β power (data now shown) from PRE to POST. A whole-brain source analysis for θ power revealed a significant cluster in left cerebellar crus I (t_21_ = 3.2, p = 0.002, Fig. 5B), showing an increase from PRE to POST. There were no correlations between POST-PRE θ power in either cerebellar or cortical locations and the angular error differences EA1-BL (p > 0.7).

**Figure 5.**
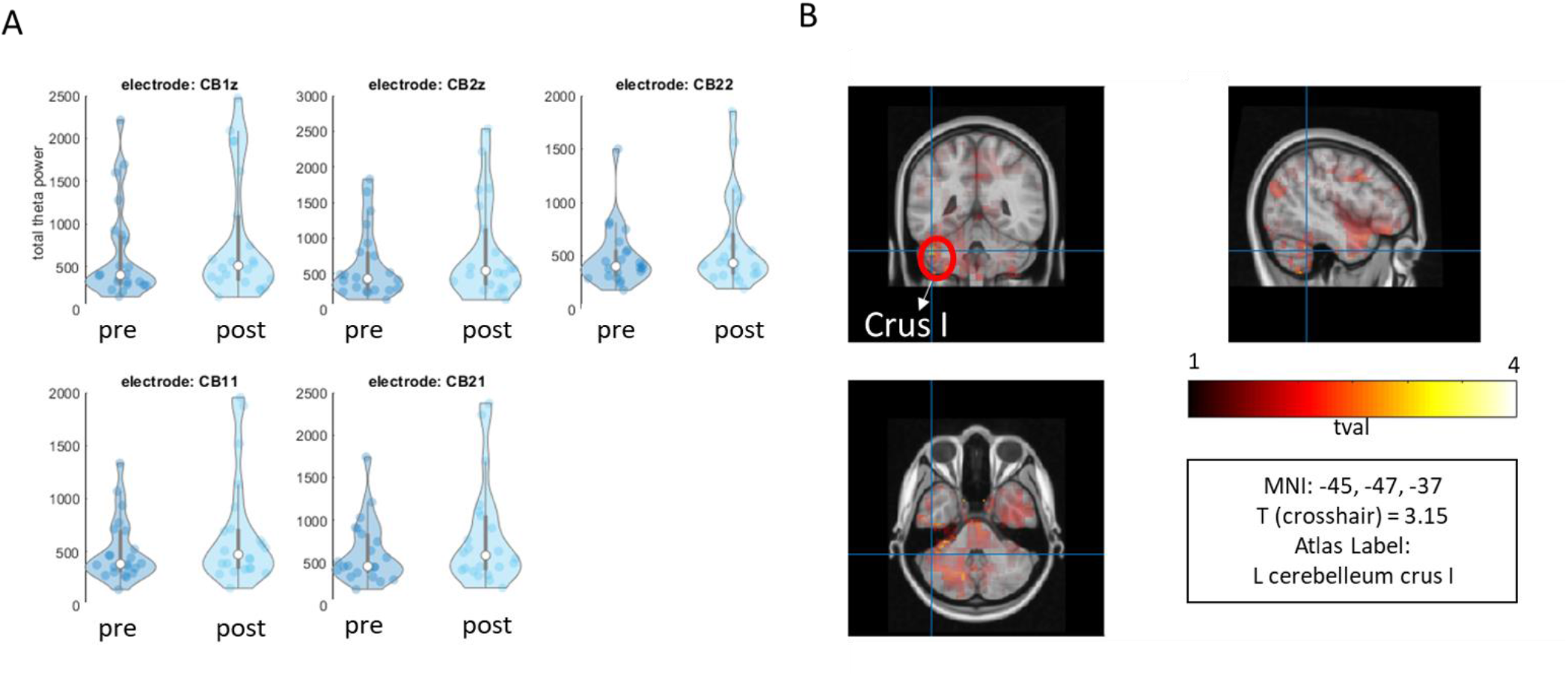
Resting state theta oscillations. **A** Theta power increased from pre-task RS to post-task RS in electrodes over the cerebellum. **B** source analysis localized the effect to left cerebellar crus I.

### 2.5. Modulation of cerebellar – SMA connectivity by visuomotor adaptation

To address the question whether interactions within the cortico-cerebellar loop are mediated by θ oscillations, we employed the phase-slope index (PSI) and studied directed changes in connectivity between left cerebellar lobule VI (lCB) and right SMA (rSMA) ROIs. The signals were extracted around the peak activation found above (see Fig. 3C and Fig. 4C). In addition, to control whether the effects are specific to the cerebellum and are not merely a reflection of occipital θ oscillations, we also analyzed connectivity between left occipital lobe (lOcc) and rSMA ROIs. Comparing EA1 to LA2, we found an increase in the directed interaction rSMA –> lCB (Z = 2.6, p = 0.009, Fig. 6). No other differences between conditions were observed in rSMA –> lCB connectivity, however connectivity tended to increase from EA1 to BL (Z = 1.8, p = 0.07, Fig. 6). Differences in directed connectivity lOcc –> rSMA between conditions were not significant (p > 0.1). There were no correlations between changes in angular error and rSMA –> lCB connectivity. Connectivity between lCB and rSMA during RS did not differ between PRE and POST (p > 0.1).

**Figure 6.**
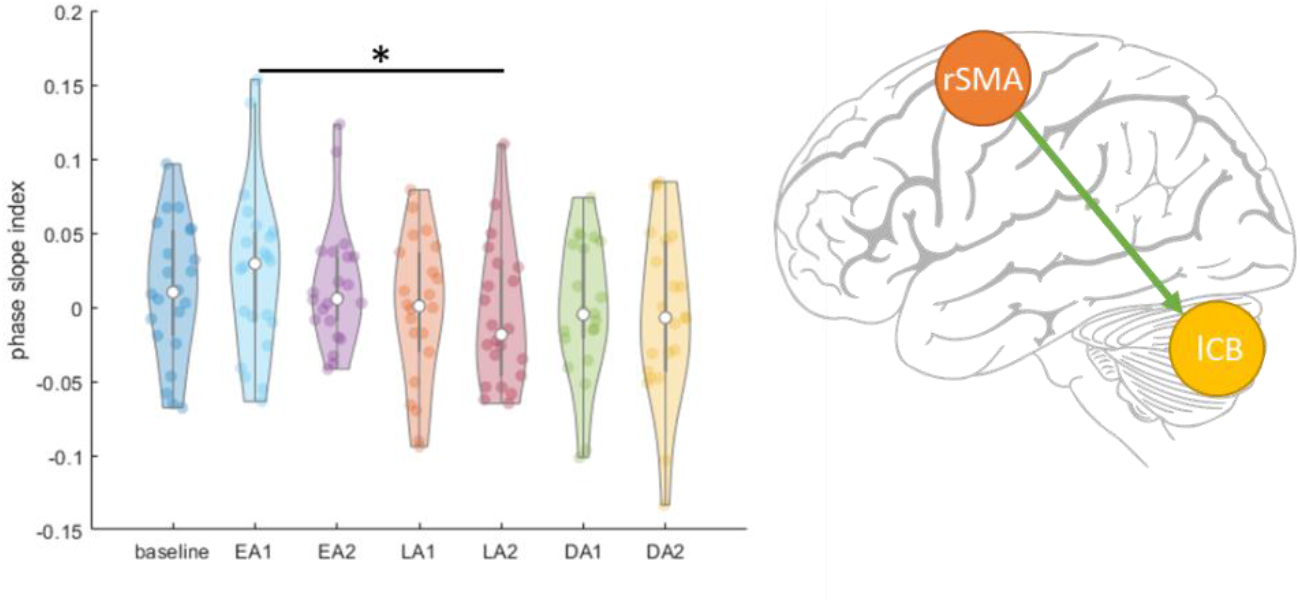
Connectivity between left cerebellum crus I and right SMA. Early adaptation showed significantly stronger connectivity expressed as the phase-slope-index, compared to late adaptation.

## Discussion

In this study, we investigated oscillations as a neurophysiological marker for communication within visuomotor adaptation networks. We found evidence for changes in theta (θ, 4-8 Hz) power with adaptation to a visual perturbation in two distinct electrode-space clusters: a central-cluster and a cerebellar-cluster. In both clusters, θ power increased with first exposure to the perturbation, attenuated with time, and rebounded when the perturbation was removed. Modulation of θ power across different adaptation phases directly followed activation patterns observed in our previous fMRI study using the exact same task (Fig. 3D, Tzvi et al., 2020). Importantly, θ power differences were directly linked to changes in the angular error: subjects who better adapted to the visuomotor perturbation early-on, had larger θ power differences (compared to baseline) in electrodes placed over the left cerebellum (CB11, CB21) as well as electrode C4. Localization of θ power changes with adaptation using individual anatomical images suggested that sources for this effect reside in left cerebellar crus I (lCB) and right SMA (rSMA). Finally, we addressed the hypothesis that cortico-cerebellar interactions are mediated by θ oscillations by analyzing the phase-slope index (PSI) between rSMA and lCB during adaptation. We found increased rSMA–>lCB interaction during early-adaptation compared to late adaptation, suggesting that θ oscillations underlie information transfer in the cortico-cerebellar network, essential for visuomotor adaptation processes.

Communication within reverberating cortical-cerebellar loops is mainly mediated through deep cerebellar nuclei and the thalamus (Bostan et al., 2013; Nashef et al., 2018). To coordinate movement, the cerebellar cortex inhibits the deep cerebellar nuclei through Purkinje cells output, which in turn excites the thalamus. The thalamus then exerts exciting influences on the motor and pre-motor cortices. The exact mechanism underlying this communication remains unclear. One theory suggests though that oscillatory synchronization could serve as a marker for information transfer within neural networks (Fries, 2005). Accordingly, cerebellar oscillations may represent as key for communication within the cortical-cerebellar loop during visuomotor adaptation.

Evidence for both low and high-frequency cerebellar oscillations is supported by previous recordings from cerebellar granule-cell layer in animals (for an overview see: De Zeeuw et al., 2008). In particular, low-frequency cerebellar oscillations (4-25Hz) may serve to spatiotemporally organize communication through output to Purkinje cells as well as through synchronization between the cerebellum and cerebral cortex (Courtemanche et al., 2013). Interestingly, LFP recordings from medial prefrontal cortex and cerebellum in guinea pigs showed increased synchronization in 5-12Hz frequency band during eyeblink conditioning (Chen et al., 2016). Accordingly, these authors proposed that medial prefrontal cortex and cerebellar communication via θ-band synchronization contributes to adaptive associative learning behavior. Our results support and extend these findings, demonstrating interactions from right medial frontal cortex to the right cerebellum during early visuomotor adaptation in humans.

Our findings further converge with previous EEG studies in humans. Using EEG source-localization, Mehrkanoon and colleagues (2016) showed increased functional connectivity in motor cortex-cerebellar interactions in oscillations around 10Hz, during resting-state directly following training of a motor task. Importantly, they showed that the cerebellum phase-lags the motor cortex, which means that training of a motor task leads to enhanced directed information transfer from motor cortex to cerebellum. The effect of visuomotor adaptation on cortical oscillations is also well studied with findings demonstrating effects in several frequency bands, including θ. For instance, Perfetti et al. (2011) found increased θ power in right posterior parietal scalp-region, for late relative to early adaptation, prior to movement initiation. This contrasts with current findings showing decreased θ power comparing late to early adaptation. Note however that effects evident during movement preparation and actual movement are distinguishable. We found no evidence to support modulation of θ power during movement preparation. The previous study additionally reported that larger θ power during late adaptation (compared to early) was associated with a larger degree of adaptation (Perfetti et al., 2011). This converges with our observation that during movement, larger θ power in early adaptation (compared to baseline) is also associated with larger adaptation. Thus, it seems that increased θ power is indicative of successful adaptation, regardless of the specific adaptation phase. Finally, low θ (2-4Hz) oscillations were found to be increased when visuomotor prediction errors, thought to reside in the cerebellum, were present (Savoie et al., 2018).

To summarize, evidence suggests that θ oscillations may play an important role in the process of visuomotor adaptation, specifically as a mechanism for communication within a cortico-cerebellar network. Here, we found that not only cortical θ oscillations are relevant for visuomotor adaptation, but most importantly cerebellar θ oscillations. In accordance with the results by Mehrkanoon et al. (2016), we found increased directed θ coherence from rSMA to lCB during early adaptation, i.e. when prediction errors are large, compared to late adaptation when prediction errors were smaller.

Most commonly associated with learning and memory, vast empirical data suggest that θ oscillations are important for establishing temporal associations between sensory stimuli (for review see: Herweg et al., 2020). On the other hand, the cerebellum has been strongly associated with temporal representations of events, known as event timing (Ivry et al., 2002). Recently, it has been shown that the cerebello-thalamo-cortical pathway affects motor cortical firing at movement onset through synchronous bursts, which could serve as key for timing motor actions (Nashef et al., 2018). In our experiment, timely associations between proprioception and visual feedback develop as subjects adapt to the visual perturbation. Therefore, we speculate that increased cerebellar θ power reflects temporal integration of proprioception in the cerebellum (Bhanpuri et al., 2013) together with action programming in SMA. SMA might communicate the programmed action to the cerebellum when a mismatch between the action and the perception of the action on the screen occurs via a θ code, leading to increased θ connectivity during early adaptation.

The present results further converge with our previous fMRI study that investigated interactions within a cortico-striato-cerebellar network underlying visuomotor adaptation (Tzvi et al., 2020). Linear decrease in activity with adaptation was maximal in left cerebellum crus II as well as right SMA, corroborating the EEG findings here. Remarkably, θ power changes with adaptation followed the exact same activation pattern in the fMRI study: fMRI activity increased significantly when the perturbation was first introduced, decreased with adaptation and rebounded when the perturbation was removed during de-adaptation (Fig. 3D). These results suggest that the observed fMRI activation pattern likely reflect changes in theta power during visuomotor adaptation. Connectivity analysis using dynamic causal modelling revealed distinct modulation of interactions in a cerebellar-premotor cortex loop by the adaptation process. Note that both the premotor cortex as well as the SMA were evident in the fMRI analysis. Thus, the results of the current study not only replicate but also expand our previous fMRI findings in that they provide a possible mechanism in which communication with a cortico-cerebellar loop could take place. A future validation of this notion could be achieved for example using simultaneous EEG-fMRI recordings during performance in a visuomotor adaptation task.

## Conclusions

In this study we demonstrate that cerebellar θ oscillations play a critical role in the process of visuomotor adaptation. Moreover, we propose that communication between cerebellum and SMA is mediated through coherent θ oscillations during the early adaptation phase, possibly for temporal integration of proprioception information from the cerebellum to create a motor program in the SMA. These findings may motivate the use of therapeutic non-invasive electric stimulation protocols in patients with cerebellar degeneration or cerebellar stroke, who have strong deficits in adaptation to a visuomotor manipulation.

## Acknowledgments

The authors would like to thank Christoph Mühlberg and Anna Wolff for their assistance with the study design. This study is supported by the DFG grant TZ 85/1-1 to ET. We acknowledge support from the University of Leipzig for Open Access Publishing.

